# Characterization of individual HIV-1 budding event using ultra-fast atomic force microscopy reveals a multiplexed role for VPS4

**DOI:** 10.1101/2021.12.12.472262

**Authors:** Shimon Harel, Yarin Altaras, Dikla Nachmias, Noa Rotem-Dai, Inbar Segal, Natalie Elia, Itay Rousso

## Abstract

The assembly and budding of newly formed human immunodeficiency virus-1 (HIV-1) particles occur at the plasma membrane of infected cells. Whereas the molecular basis for viral budding has been studied extensively, investigation of its spatiotemporal characteristics has been limited by the small dimensions (< 100 nm) of HIV particles and the fast kinetics of the process (a few minutes from bud formation to virion release). Here we applied ultra-fast atomic force microscopy to achieve real-time visualization of individual HIV-1 budding events from wildtype (WT) cell lines as well as from mutated lines lacking vacuolar protein sorting-4 (VPS4) or visceral adipose tissue-1 protein (VTA1). Using single-particle analysis, we show that HIV-1 bud formation follows two kinetic pathways (fast and slow) with each composed of three distinct phases (growth, stationary, decay). Notably, approximately 30% of events did not result in viral release and were characterized by the formation of short (rather than tall) particles that slowly decayed back into the cell membrane. These non-productive events became more abundant in VPS4 knockout cell lines. Strikingly, the absence of VPS4B, rather than VPS4A, increased the production of short viral particles, suggesting a role for VPS4B in earlier stages of HIV-1 budding than traditionally thought.

## Introduction

Budding is considered a fundamental step in the replication of retroviruses, such as human immunodeficiency virus (HIV). During budding, which occurs at the surface of the host cell, the membrane is deformed outward by the assembled virus to produce a spherical particle. Several studies have suggested that Gag, which is a major structural protein, has an intrinsic curvature that produces spherical particles during in vitro assembly ^1–5^. Indeed, electron microscopy analysis of budding virions revealed spherical Gag structures beneath the cell membrane ^6^. Nonetheless, current understanding of the mechanism underlying retroviral budding is quite limited because of the considerable complexity associated with measuring real-time bud formation.

Increasing evidence suggests that cellular factors play an important role in the budding of viruses from the cell membrane. In several viruses, including equine infectious anemia virus (EIAV) ^7^, vaccinia virus ^8^, and HIV ^9–12^, actin was suggested to participate in the budding process and a few viruses were shown to remodel the actin cytoskeleton of their host ^13, 14^. In addition, studies have shown that HIV-1 exploits the cellular machinery of the endosomal sorting complexes required for transport (ESCRT) to accomplish particle release ^15^. Given the involvement of host cellular machineries in viral budding, gaining a realistic understanding of the process requires its study in the context of whole live cells.

The ESCRT machinery plays a significant role in several cellular processes, such as multivesicular body (MVB) biogenesis ^16^ and cytokinetic abscission ^17^. Comprised of five subcomplexes (ESCRT-0, ESCRT-I, ESCRT-II, ESCRT-III, and VPS4-VTA1), ESCRT machinery facilitates the membrane remodeling processes required for budding away from the cytoplasm. In HIV budding, the recruitment of early ESCRTs (ESCRT-I, ESCRT-II) ^18, 19^ and ALG-2 interacting protein X (ALIX) ^20–22^ is mediated by the p6 domain located at the C terminal end of Gag and facilitates the recruitment of ESCRT-III to the bud’s neck. ESCRT-III then recruits the final ESCRT subcomplex, namely the VPS4–VTA1 complex formed by vacuolar protein sorting-4 and visceral adipose tissue-1, with this complex required for the scission of the viral bud’s neck from the membrane during the final stages of budding and involved in the disassembly and recycling of the ESCRT-III complex ^15, 19, 23, 24^.

Advances in quantitative fluorescence single particle imaging allowed researchers to obtain a detailed description of the viral budding process. Using these approaches, the dynamics of HIV biogenesis and budding ^25, 26^ and, more recently, the recruitment of ESCRT proteins during budding were described ^27, 28^. Although these studies provided unique information on the sequence of protein recruitment during the budding process, they relied on fluorescent tagging of the proteins, which is prone to affecting the budding process, ^29^ and were limited to budding stages associated with changes in fluorophore density.

The entire process of viral budding can be visualized using atomic force microscopy (AFM), as we have previously demonstrated for budding of murine leukemia viruses (MLVs) ^30^. Operating the AFM in the torsional mode, we were able to track dynamic changes in local actin during MLV and HIV budding ^14^. Although the temporal resolution of standard AFM (~4–5 min/image) is inferior to that of total internal reflection fluorescence microscopy (TIRFM), it is sufficient for the purpose of monitoring MLV budding, which occurs on a time scale of a few tens of minutes ^30^. However, HIV budding is characterized by fast kinetics (on a time-scale of only a few minutes, based on fluorescence microscopy experiments) ^26, 31^ and consequently it is not possible to obtain a detailed characterization of the kinetics of an individual HIV budding event using standard AFM ^14^. Recent advances in force microscopy have dramatically improved its temporal resolution, enabling the acquisition of AFM images every few seconds. Such a temporal resolution combined with high resolution in the z-axis allows the nascent bud to be tracked from its first appearance until it is released into the surrounding media, and beyond. In this study, we applied the fast AFM approach to describe the spatiotemporal characteristics of HIV-1 budding kinetics by tracking individual budding events in live human cells (HeLa). Using the clustered regularly interspaced short palindromic repeats-Cas9 (CRISPR-Cas9) gene editing method to knock out ESCRT proteins VPS4A, VPS4B, and VTA1, we further characterized the contribution of the VPS4-VTA1 complex to viral budding.

## Results

Budding of HIV-1 particles was imaged with the AFM operated in tapping mode to minimize possible damage to the cells by the probe. Topographical images of non-transfected HeLa cells revealed a relatively smooth membrane surface (Figure 1A). In contrast, the surface of HeLa cells infected with HIV-1 was covered by protrusions of different sizes, which were likely virus particles in various budding phases (Figure 1B). Due to convolution between the AFM probe and particles, the width of the particles often appeared to be larger than their expected native size (110–140 nm) ^32^ and therefore we measured particle height ^33^. In addition, we used phase imaging to support the detection of viral particles. During intermittent mode imaging, tip-surface interactions cause energy dissipation that produces a shift between the scanner and the tip amplitudes. This phase shift was used to track changes in the surface stiffness ^34, 35^ and thereby rule out possible non-viral protrusions made by variations in membrane curvature. Combining the height and phase data allowed us to detect budding from very early in the process.

**Figure 1:**
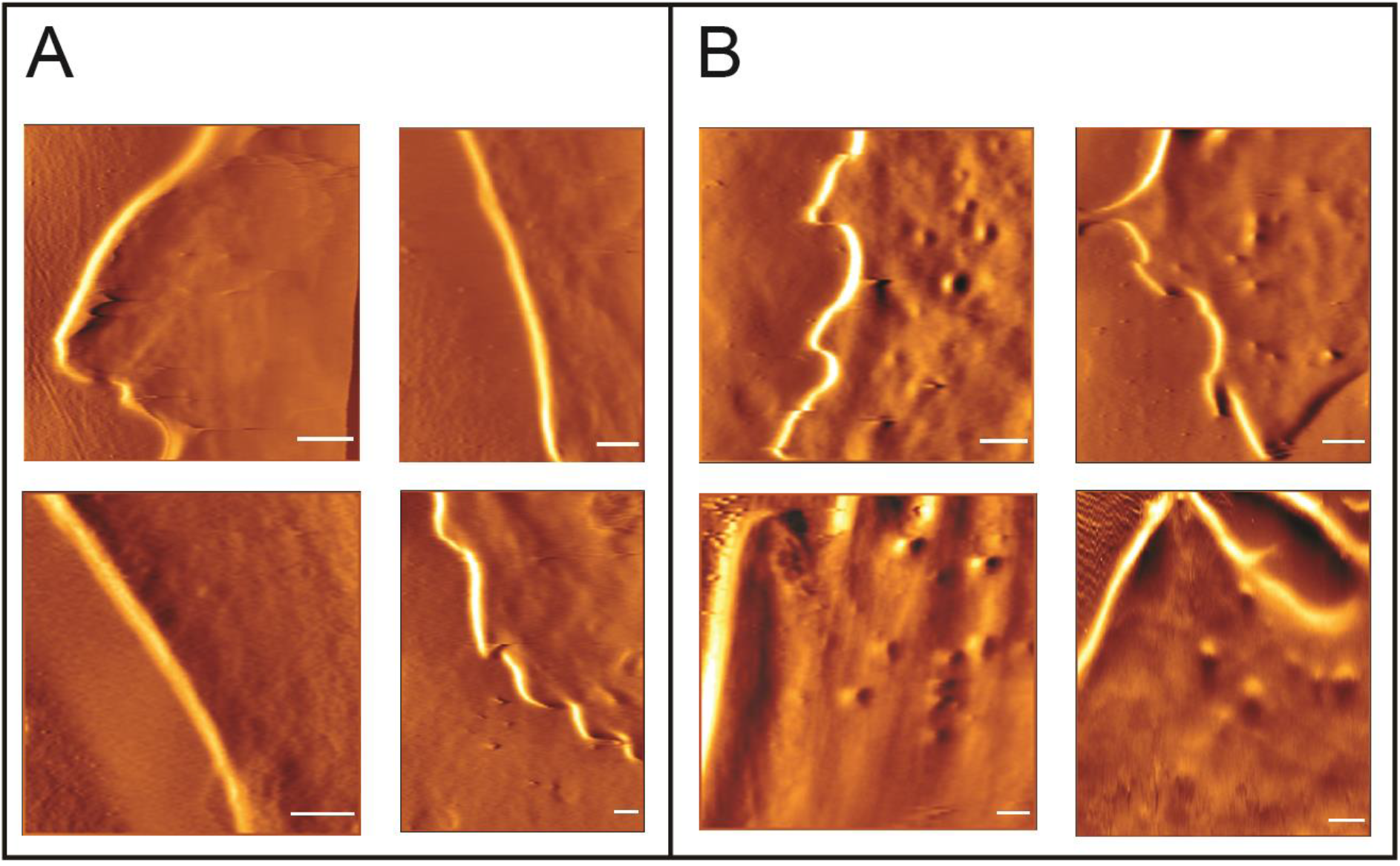
AFM images of uninfected and HIV-1 infected wildtype HeLa cells. (A) Typical AFM amplitude images, acquired in intermittent mode, of uninfected live cells showing the relatively smooth cell membrane. (B) AFM amplitude images of HIV-1 infected cells. Virus particles at different stages of budding appear as protrusions of different heights on the cell membrane. Due to convolution between the AFM tip and the virus, these protrusions appear larger than their actual dimensions. Scale bars in all images are 1 μm.

To characterize the kinetics of HIV-1 budding, single particles were analyzed, frame by frame, from their initial appearance on the cell surface until their complete disappearance (Figure 2A). Consistently with previous reports that used AFM to capture budding events ^14, 30^, we usually visualized such events on the surface of the cell with the direction of budding parallel to the z-axis. In addition, we captured several budding events from the side of the cell (Figure 2B) to image bud neck formation. We analyzed a total of 89 budding events captured from 32 WT-HeLa cells.

**Figure 2:**
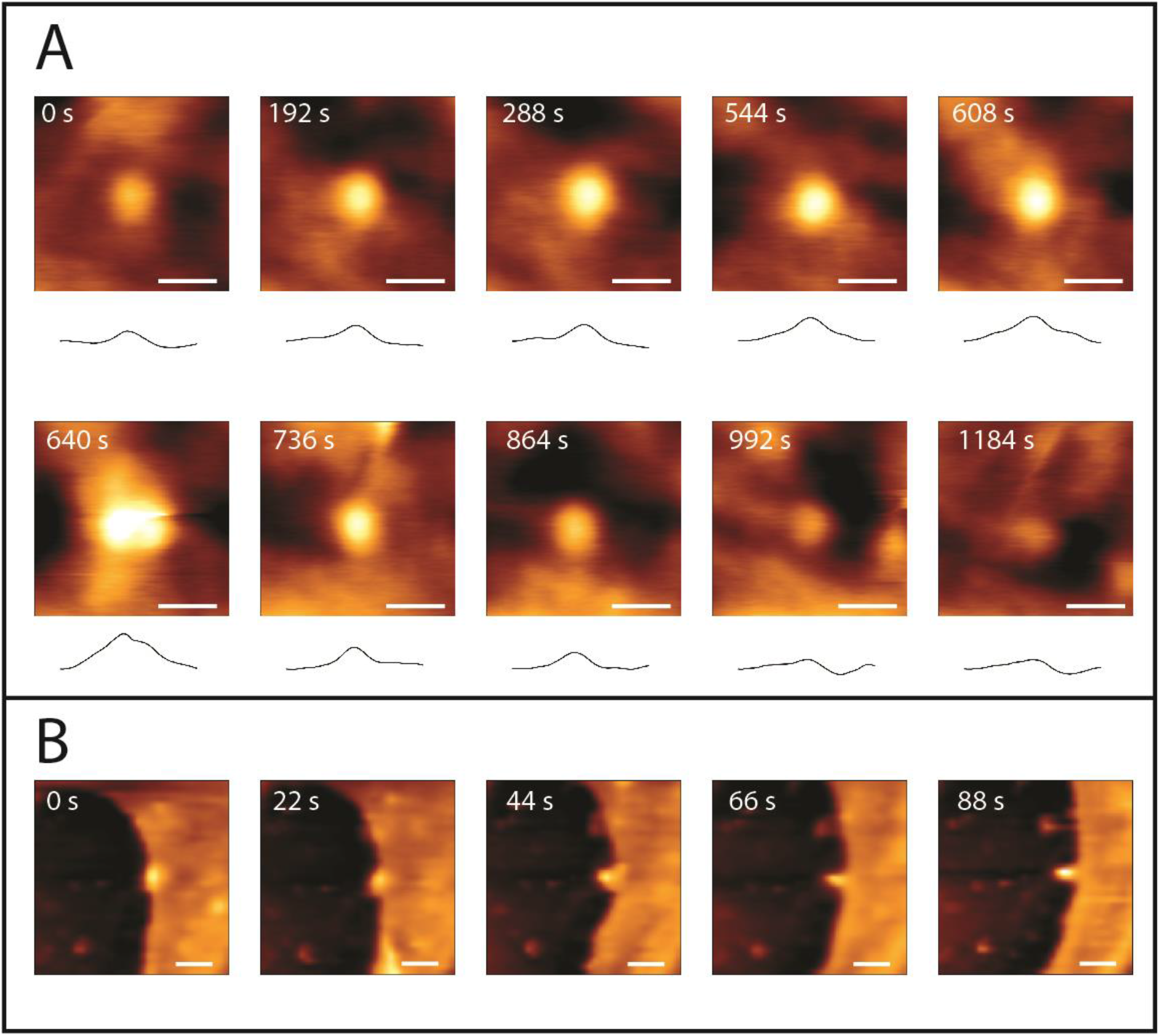
Visualization of a virion budding event. (A) Topographic imaging taken using fast AFM showing the complete budding sequence, from initial virion assembly through to the release of a single virus particle from a live wildtype HeLa cell, with their corresponding cross-sectional profiles sketched below each image. At the beginning of the process the bud is seen as a small bright protrusion. After ~600 seconds, the budding virion has grown to its maximal height and it is then released from the cell membrane. Image scan area is 1.6 μm × 1.6 μm, 128 pixels × 128 pixels. Scale bar is 0.5 μm. (B) Documenting an HIV-1 budding event from the side of the cell. Scan area 2.7 μm × 2.7 μm, 128 pixels × 128 pixels. Scale bar is 0.5 μm.

The maximal height of the measured budding particles (70 ± 2.46 nm) was shorter than their expected height (110–140 nm) determined by analyzing isolated native virions ^32^. The observed reduced particle heights were likely an outcome of the force exerted by the AFM probe ^36^ pressing the virion particles into the soft membrane of the live cells. Particles were divided into two groups according to their maximal height. A threshold height value of 45 nm was determined from the particle height distribution to divide the particles into two discrete height populations: short and tall. The short group (height ≤ 45 nm) constituted 33% (29 events) of all particles and had an average maximal height of approximately 35 ± 1.47 nm. The tall group (height > 45 nm) constituted 67% (60 events) of all particles, had an average maximal height of approximately 70 ± 2.46 nm, and likely represented fully assembled virions.

We next analyzed the spatiotemporal kinetics of individual budding events (particle height vs. time) for particles from the tall (Figure 3B) and short (Figure 3C) groups (Figure 3B–C). We preformed the analyses separately for each height group because the average maximal height of the short particle group was too small to represent a fully assembled virus, and therefore their kinetic profiles were unlikely to represent a viable budding event. We divided each trajectory into three phases: 1) growth; 2) stationary; and 3) decay. The growth phase refers to the assembly of the viral particle and describes the period from the initial appearance of the particle on the membrane until it attains its maximal height. The stationary phase is the period spent by the assembled particle at its maximal height. The decay phase describes the period during which the particle is either released or (if budding was non-productive) reabsorbed, with the bud-remanent (or the non-productive bud) then declining from its maximal height until it completely disappears from the cell surface. The budding time of a particle is defined as the total period from the beginning of the growth phase until the end of the decay phase. In some cases, only minimal lifetime values could be estimated because partial trajectories were captured that covered only the first two phases (growth and stationary) or the last two phases (stationary and decay).

**Figure 3:**
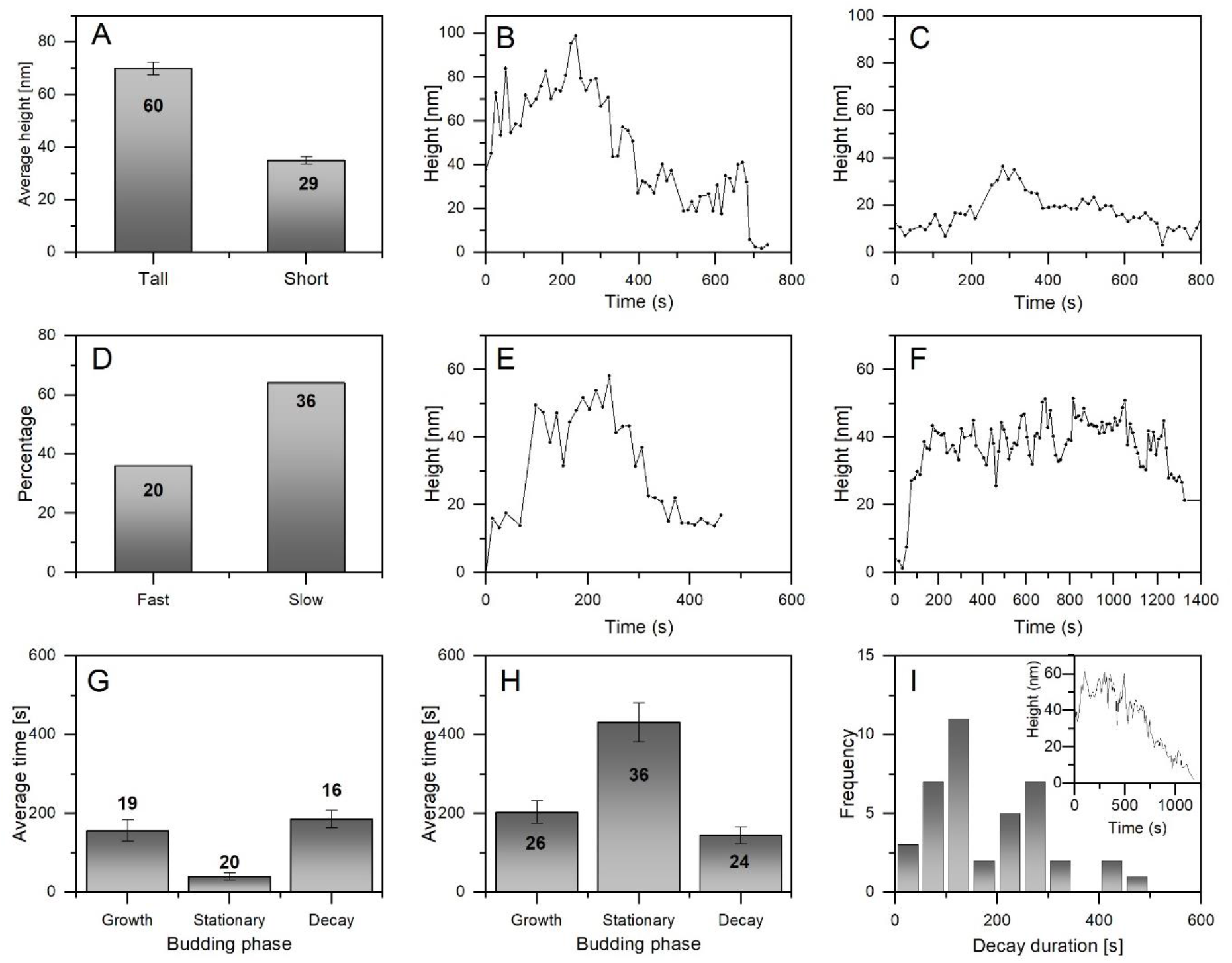
Analysis of the kinetics of single HIV-1 budding events from wildtype HeLa cells. (A) All analyzed budding events (89 in total) were divided into two height groups (short and tall) according to their maximal height, with the threshold set at 45 nm based on particle distribution. Most budding events (*n*=60, 67%) arose from tall particles, which exhibited an average maximal height of 70 nm. The remaining budding events (*n*=29, 33%) arose from short particles, which exhibited an average maximal height of 35 nm. (B–C) Representative kinetic trajectories of particles from the tall (B) and short (C) particle height populations. (D) The tall particles for which we obtained full kinetic trajectories (n=56 of the 60 relevant budding events) were further divided into two main groups according to their kinetics profiles: 20 (36%) of the tall budding events occurred without a significant stationary phase, leading to fast budding kinetics lasting for ~380 s, on average. The remaining 36 (64%) of tall budding events were characterized by slow kinetics, having an averaged lifetime of ~660 seconds. The total number of fast and slow events is less than the total number of events involving tall buds because of a few incomplete kinetics trajectories. (E–F) Representative fast (E) and slow (F) budding events. (G–H) Average durations of the three kinetic phases (growth, stationary, and decay) for fast (G) and slow (H) budding events. (I) Distribution of the durations of the decay phases in budding events involving tall particles, clearly showing two kinetics populations. The inset shows a representative kinetic trajectory of an event with a long decay rate.

The tall particle group was further divided into two budding kinetics groups: fast and slow (Figure 3D). Both groups exhibited similar growth and decay times, but the slow group had a longer stationary phase that consistently exceeded 120 s in duration and often lasted several minutes (Figure 3G, H). A short stationary phase indicated very rapid release of the virus. Among tall particles, the majority of budding events (36 events, 64%) incorporated an extended stationary phase (average time, 431 ± 50 s) and exhibited slow trajectories with a duration of 665 ± 92 s on average. A representative kinetic trajectory for a slow budding event is shown in Figure 3F. In the remaining events (20 events, 36%), the stationary phase was very brief (< 120 s), which gave rise to faster budding kinetics (Figure 3E shows a typical fast kinetic trajectory). The average total duration of the rapid budding events was 382 ± 46 s, within which the decay phase lasted an average of 186 ± 22 s. Analysis of individual particles (Figure 3I) revealed that the relatively long average decay time is somewhat misleading, as averaging flattens differences between two groups, one comprised of rapidly-decaying particles (over about 150 s) and another comprised of slow budding events comprised of slowly-decaying particles (over about 500 s; Figure 3I inset). Analysis of the budding kinetics of the short particles is more challenging since it is nearly impossible to distinguish between their stationary and decay phases because of the relatively poor signal to noise ratio. Nonetheless, the overall average decay time of the short particles (148 ± 23 s) was similar to that of the slow budding events of the tall particles (145 ± 21 s).

In several instances we observed structural features pre- or post-budding. When viewing frames from immediately before a particle was detected (*i.e*., before the beginning of the growth phase), we occasionally observed a slight preliminary bulge that usually lasted about 200 s before a clearly-defined particle was visible and the growth phase commenced (Figures 4A). Similarly, in the case of rapid decay, structural features with heights (~20 nm) that were slightly greater than the topography of the surrounding cell membrane often remained visible at the budding site for 100–1000 s post virion release (Figure 4B). Often it was difficult to distinguish these residual structures from particles with long decay phases. In several cases (~9), we observed the formation of an initial slight preliminary bulge that entered the growth phase but failed to release a virion and was then reabsorbed before being followed by a subsequent budding event that proceeded to virion release (Figure 4C).

**Figure 4:**
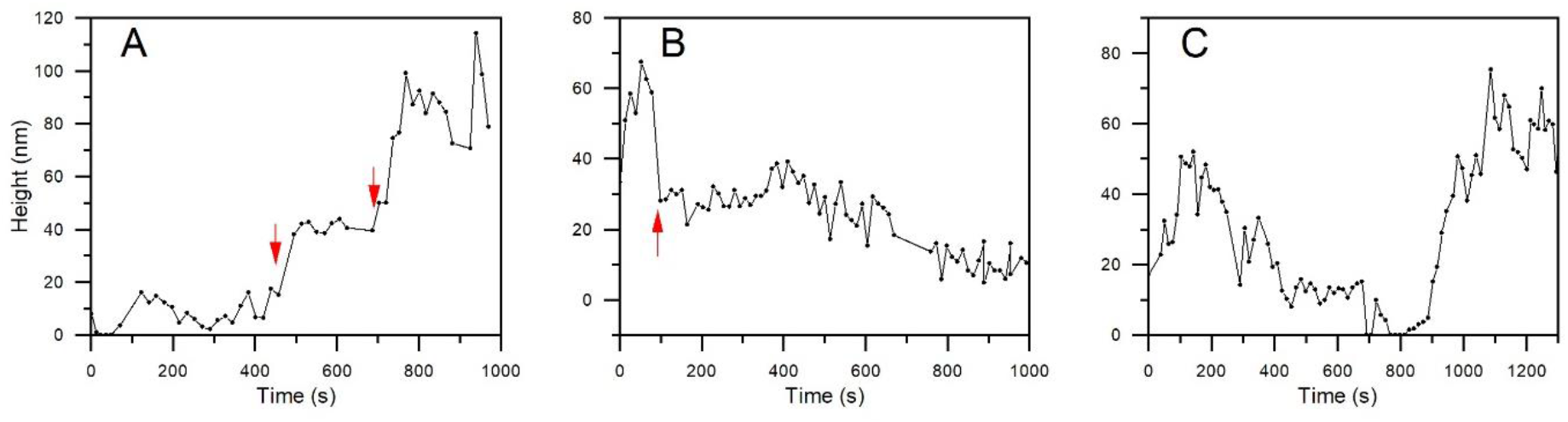
Pre- or post-budding structural features. (A) Kinetic trajectory of an event in which a slight bulge appeared prior to the growth phase. The lifetime of the pre-budding structure (labeled by two red arrows) was approximately 200 s. (B) An example of a post-budding structural feature that remained for ~800 s after the virion particle was released. The estimated beginning of the post-budding feature following the end of the decay phase is marked by a red arrow. (C) Two consecutive budding events at the same site. An initial budding event produced a slight bulge that decayed without releasing a virion and was followed by the formation of a tall particle and virion release.

Having obtained a quantitative characterization of viral budding, we endeavored to examine the contribution of the VPS4–VTA1 complex to budding height and kinetic profile. To this end we generated complete knockout (KO) cells for the VPS4 paralogs (VPS4A.KO and VPS4B.KO) and VTA1 (VTA1.KO) using CRISPR-Cas9 technology (Figure 5A). All attempts to use the CRISPR-Cas9 method to generate cells depleted of both VPS4A and VPS4B failed, suggesting that at least one VPS4 isoform is needed for cell viability. To test the effect of VPS4A or VPS4B on the overall amount of viral production, infected cell supernatants were immunoblotted using an antibody against viral capsid protein p24 (Figure 5B). Interestingly, although all three knockouts reduced virus production compared with WT cells, viral production was reduced to a much greater extent by VPS4B.KO and VTA1.KO compared with VPS4A.KO. We then used AFM to image 70, 86, and 84 budding events in HIV-1-infected VTA1.KO, VPS4B.KO and VPS4A.KO cells, respectively (Figure 5C) and compared them with the images obtained from non-infected cells (Figure 1A). Budding trajectories were acquired and analyzed using a method similar to the approach we used for virus particles produced from WT cells. Imaging of non-HIV-1 infected cells showed a relatively smooth membrane surface lacking the protrusions observed in infected cells. Using the same criteria as were used for WT cells, we divided budding particles into two groups (tall and short) according to their maximal height (Figure 5D). We found that the percentage of short particles in VPS4A.KO (37%) was marginally greater than observed in WT cells (33%). However, the percentage of short particles was significantly greater, reaching a value of almost 50% in VPS4B.KO cells (41 events, 48%) and VTA1.KO cells (34 events, 49%).

**Figure 5:**
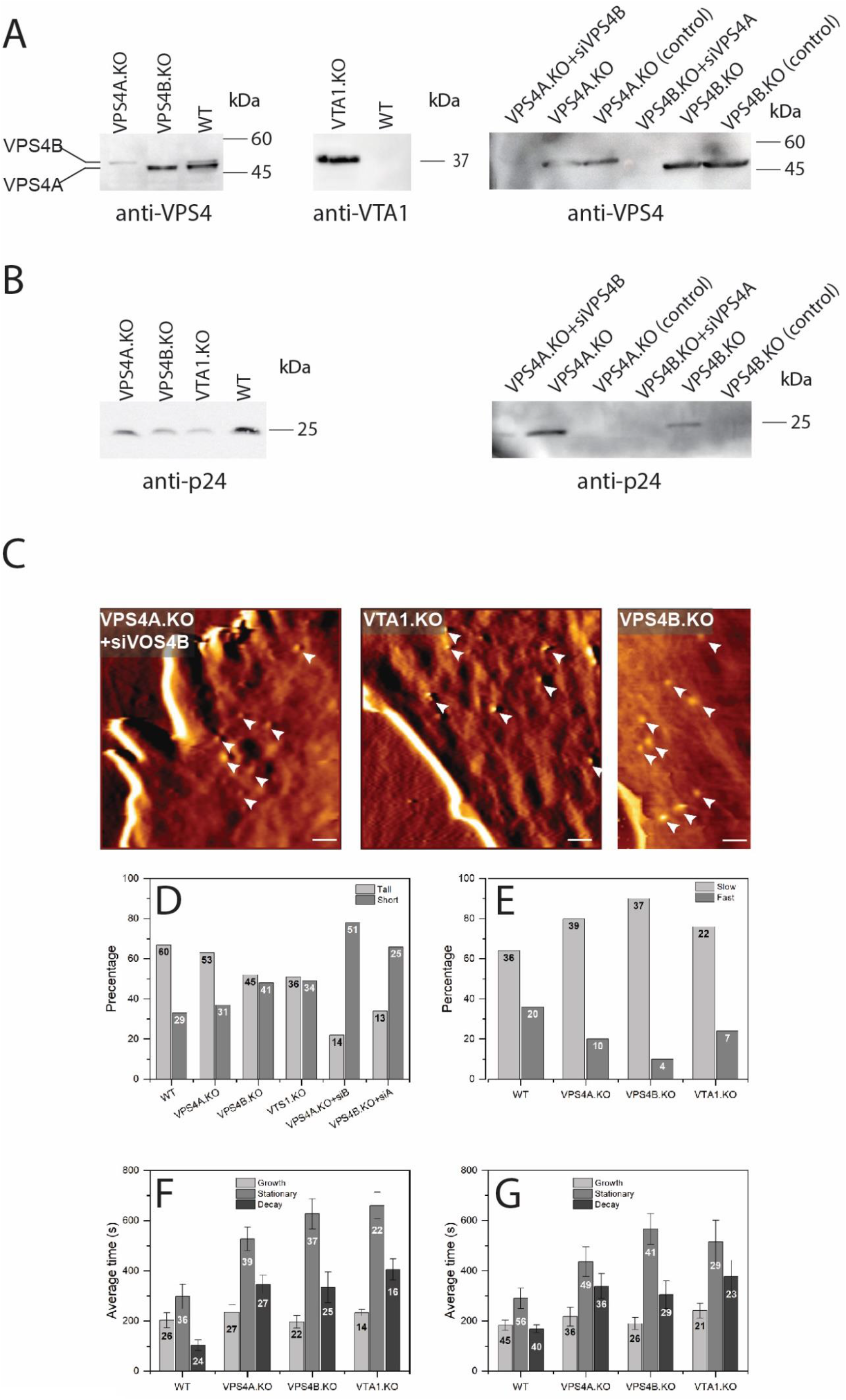
Analysis of the kinetics of single HIV-1 budding events in VPS4 and VTA1 knockout HeLa cells (VPS4A.KO, VPS4B.KO, and VTA1.KO) compared with wildtype (WT) HeLa cells. (A) Verifying VPS4/VTA1 depletion. WT or KO HeLa cells were directly lysed (left and middle panels) or transfected with an appropriate siRNA (siVps4B for VPS4A; siVps4A for VPS4B) and then lysed (right panel). Next, equal amounts of the lysed cells were loaded on SDS-PAGE and subjected to western blot analysis using the antibodies specified beneath each panel. (B) Testing viral production. The supernatants were collected from WT and KO HeLa cells transfected or not with siRNA probes, as indicated, and infected with HIV-1. Normalized amounts of supernatant were loaded on SDS-PAGE and subjected to western blot analysis using antibodies for the viral capsid protein p24. (C) Representative AFM images of cells depleted of different VPS4-related proteins, as indicated, and infected with HIV-1 particles. Budding events are marked with white arrows. Nearly all budding events had a low maximal height (*i.e*., produced short, as opposed to tall, buds). The average maximal heights for the labeled events were 10 nm, 27 nm, and 13 nm, respectively. Scale bar is 1 μm. (D–E) The relative populations of budding events producing tall and short particles (D) and having slow and fast kinetics (E) shown as a percentage (y-axis) and as the number of events (given on each bar). Average time spent in each of the three kinetic phases, growth, stationary, and decay, with the number of slow (F) and total (G) budding events shown on each bar. Error bars indicate SEM.

We next sought to study the effect of deficiency in both VPS4 paralogs on HIV-1 budding. To overcome the challenge of cell viability, we used siRNA to reduce the level of one VPS4 isoform in KO cells that were already depleted in the other isoform. Figure 5A (right panel) shows that using combining these techniques enabled us to reduce significantly the levels of both VPS4 variants in the same cells. Strikingly, we observed a significant increase in the population of short particles in both VPS4A.KO+siVPS4B (78%) and VPS4B.KO+siVPS4A (66%) cells, which was accompanied by the almost complete arrest of virus production, as detected by p24 antibodies (Figure 5B right panel, 5D).

Finally, we analyzed the kinetics of viral budding in VTA1.KO, VPS4B.KO, and VPS4A.KO cells. Figure 5E shows that the percentage of slow budding trajectories was increased in VPS4A.KO (80%), VPS4B.KO (90%), and VTA1.KO (76%) compared with WT (64%). We found that these slow budding event trajectories (Figure 5F) had a considerably longer stationary phase (594 ± 53 s) and significantly longer decay phase (355 ± 47 s) compared with WT (stationary, 298 ± 60 s; decay, 103 ± 21 s). As demonstrated in Figure 5G, the lengthening of these phases in the slow events has retarding effect on the overall kinetics of budding.

## Discussion

We have previously utilized AFM to analyze the budding kinetics of MLV and HIV-1 particles ^14, 30^. In the earlier studies, the low temporal resolution of standard AFM (~ 4 min/frame) was sufficient to characterize the relatively slow kinetics of MLV budding but was insufficient to provide a comprehensive examination of HIV-1 budding. In the current study, live cells were imaged repetitively using ultra-fast AFM over the course of several hours, during which HIV-1 budding events from the cell’s membrane were recorded. This AFM setup can acquire images at a temporal resolution of 4–8 s/frame. Combining the high topographical resolution of AFM with a newly-available high temporal resolution provided a step-by-step description of individual HIV-1 budding events.

Our analysis revealed considerable variability in HIV-1 budding times. We found that the duration of an individual HIV-1 budding event was 150–1500 s, with an average duration of roughly 532 ± 58 s (~8.9 min). These values are in agreement with previous studies that used TIRFM ^25, 26, 31^. We defined two kinetic pathways for HIV-1 budding: fast and slow. We found that both pathways had similar growth and decay phases. However, between the growth and release phases, budding kinetics trajectories from the slow pathway included a significantly longer stationary phase (characterized by constant particle height) than was found in the fast pathway. Interestingly, despite the budding kinetics being significantly slower for MLV compared with HIV-1, similar slow and fast pathways were observed ^30^. The similarity between MLV and HIV-1 may indicate a shared mechanism for the assembly and budding of enveloped retroviruses. At this stage, we cannot conclusively determine the mechanism underlying the observed slow and fast budding pathways. Jouvenet et al. ^26, 31^ proposed that reduced accessibility to cytosolic Gag slows the kinetics of budding. In all of our measurements, the virion particles were found to have very similar growth rates, regardless of their overall kinetics. Furthermore, the stationary phase appeared after the virion had reached its full or nearly full size for all analyzed particles. Therefore, we propose that the slow pathway and specifically the stationary phase are associated with scission of the viral bud’s neck from the membrane, whereas Gag recruitment is not associated with the fission step but rather occurs during the growth phase.

We further found that budding events can be divided into two groups according to maximal particle height. The short group, comprising budding particles with a low maximal height, is of particular interest as these buds are too short to represent fully assembled virions and have a decay phase that is too long to represent the release of a virion from the cell surface. It should be noted, however, that similar slow-decaying budding events were also observed for some of the tall budding particles. Our VPS4 knockout experiment showed that, in cells depleted both VPS4A and VPS4B, nearly all budding events involved short particles. Intriguingly, virus production was severely impaired in these cell lines, according to western blot analysis. We therefore propose that these slow budding events that produce short particles represent a non-productive pathway that produces partially or incorrectly assembled virions. Ultimately, this budding pathway does not lead to successful particle release, with its products being slowly absorbed back into the cell. Reverse budding trajectories have been proposed previously for HIV-1 budding ^26, 31, 37, 38^. In these studies, the newly formed viral particles were proposed to undergo internalization through the endosomal pathway. Alternatively, these non-productive budding events may be due to insufficient recruitment of the ESCRT machinery proteins to budding sites, as has been observed previously in the budding of MVBs ^39–41^.

The functional differences between VPS4A and VPS4B with respect to viral budding are currently unknown. To address this question, we measured HIV-1 budding in knockout cell lines lacking one VPS4 paralog. We found that the removal of either VPS4A or VPS4B had analogous effects on the kinetics of HIV-1 budding. In both cases, the percentage of fast budding events significantly decreased compared with WT. A detailed analysis of the kinetics trajectories revealed that the stationary and decay phases were nearly two-fold longer when either VPS4A or VPS4B was absent, compared with WT. However, removing VPS4B significantly increased the percentage of budding events producing short particles, and dramatically impaired virus production. In contrast, in VPS4A.KO cells, virus production (based on p24) and particle height distributions were similar to those found for WT cells, suggesting that VPS4B is the more potent isoform in viral budding. These findings are consistent with previous data showing that expressing an siRNA-resistant VPS4B construct was able to rescue viral release in cells in which both VPS4A and VPS4B had been deleted using siRNA^42^. To the best of our knowledge this is the first direct demonstration of a functional difference between the two paralogs of VPS4 in HIV-1 budding.

VTA1 is known to stabilize the hexameric, active conformation of VPS4 and to increase its ATPase activity ^43^. We therefore expected that VTA1.KO cells would exhibit a phenotype similar to that of a VPS4A+B double KO. However, this was not the case. Although cells in which CRISPR and siRNA had been used to achieve depletion of both VPS4A and VPS4B indeed exhibited the most severe phenotype in terms of slow viral production (p24) and short particle height, VTA1.KO cells exhibited a less severe phenotype that was similar to that observed in VPS4B.KO cells. These results suggest that, while both VPS4A and VPS4B participate in viral budding, VTA1 primarily affects the function of the VPS4B isoform, at least in the context of viral budding. It will be interesting to examine whether this difference also applies to the function of the two VPS4 isoforms in other cellular processes and whether, in vitro, VTA1 affects VPS4A and VPS4B hexamerization and ATPase activity to different extents.

The observed effect of VPS4 on virus production is in agreement with the current view that the cell’s ESCRT machinery is recruited in the late stages of viral budding and plays a crucial role in the fission of the newly formed bud ^19, 28, 31^. However, the findings that, in the absence of VPS4 and specifically in the absence of VPS4B, particle height is significantly reduced suggest a role for VPS4 in earlier budding stages, such as during the growth phase. These observations are in line with previous studies suggesting VPS4 is required both for proceeding from early to late budding stages and for final scission during MVB formation ^39^.

## Materials and Methods

### Cell culture and transfection

For cell culture, naïve and mutated HeLa cells were cultured in Dulbecco’s modified Eagle medium (supplemented with 10% heat-inactivated bovine serum, 1% penicillin-streptomycin, and 1% glutamine) and incubated at 37 °C in 5% CO_2_. For AFM imaging, cells were plated in 35 mm cell culture dishes and grown to ~50% confluence. To produce HIV-expressing cells, HeLa cells were transfected with ΔEnv IN-HIV-1 plasmid (DHIV3-GFP-D116G). For siRNA transfection, siRNA target sequences were designed using Integrated DNA Technology (siVPS4A: CCGAGAA GCUGAAGGAUUAUUUACG and siVPS4B: GUACAGUCAGCUACUCAUUUUAAAA). Transfections were performed using lipofectamine 2000 or jetPRIME® (TAMAR) transfection reagents. Cells were co-transfected with green fluorescent protein (GFP) to identify HIV-1 transfected cells using fluorescence microscopy. AFM imaging was typically performed 24–36 h after transfection. The culture medium was replaced with fresh medium before transfection and imaging.

### Knockout cell lines

VPS4A.KO or VPS4B.KO HeLa cells: for CRISPR/Cas9-mediated gene disruption, sgRNAs for VPS4A (AGTGCGTGCAGTACCTAGAC) or for VPS4B (CAAACAGAAAGCGATAGATC) were cloned to the lentiCRISPR plasmid (Addgene, #49535). Following transfection and puromycin selection, single clones were isolated and expanded. KO clones were validated by western blot analysis using anti-VPS4 antibody (Sigma, SAB4200025).

### AFM imaging of live cells

AFM imaging was carried out using a JPK Nanowizard Ultra-Speed AFM (JPK Instruments-Bruker, Berlin, Germany) mounted on an inverted optical microscope (Axio Observer, Carl Zeiss, Heidelberg, Germany). Images of cells were acquired in AFM fast mode at scanning rates of 5–30 Hz in a fluid environment. Probes designed for optical performance in fluid at high scanning speeds were used (USC-F0.3-k0.3, Nanoworld). The typical resonance frequencies of the probes in liquid were 90–150 kHz and their corresponding spring constants were 0.2–0.5 N/m. Spring constants were experimentally determined by measuring the thermal fluctuations of the cantilevers.

For budding imaging, cells transfected with HIV-1 were imaged repetitively over a period of 2–3 hours. To minimize damage to the cells by the AFM probe and allow high-speed imaging, scans were made with the AFM in intermittent scanning mode. Due to the relatively low z-axis range of the scanner, all measurements were performed near the periphery of the cells, where they are more flattened. The dimensions of the scanned areas was usually between 5×5 μm^2^ and 10×10 μm^2^. The acquisition time of a single image varied from 5–20 s, depending on the scan size, scan rate, and the number of scan lines. To maintain the health of the cells properly during measurements, the medium was replaced with fresh medium before imaging. Cells were maintained at pH 7.4 (20–25 mM HEPES buffer) at a constant temperature of 36.7 °C using a heat controller (PetriDish Heater, JPK-Bruker). Under these conditions, the cells remained healthy and viable for up to 3 hours.

### Image rendering and kinetic analysis

The kinetics of viral budding was determined by measuring the viral protrusion height as a function of time. Identification of early budding in the height channel was challenging due to their low height relative to the rough surface of the cell membrane. Therefore, virus particles were initially detected in the amplitude image, as it usually had better contrast than the corresponding height image. Once virus particles were identified, their images were rendered using JPK Data Processing software (Version 7.0.130). To determine particle height, a cross-sectional profile was extracted from each frame. The corresponding height was calculated using a script written in MATLAB software (The Math Works, Natick, MA) by subtracting the height of the local cell surface from the highest point of the particle. In this method, single budding events were tracked frame by frame to extract their kinetics profile, which was constructed from all cross-sections.

### Western blot assay

HeLa cells were lysed 24 h post-transfection using RIPA lysis buffer (150 mM NaCl, 1% Nonylphenoxypolyethoxyethanol (Tergitol NP-40), 0.5% deoxycholate, 0.1% sodium dodecyl sulfate (SDS), 50 mM Tris [pH 8.0]) supplemented with complete protease inhibitor (Cat K1010, ApexBio) for 30 min at 4 °C. Total protein concentrations were measured using the BCA Protein Assay Kit (Cat #5000206, BIORAD), and equal total protein amounts were loaded in each lane. For the virus production assay, which was conducted 24 h post transfection, the transfection medium was replaced with fresh culture medium and collected 8 hours later. Supernatants were normalized against the respective total amount of protein measured. Membranes were stained with anti-VPS4 (1:750, Sigma, SAB4200025), anti-P24 (1:10000, 183-H12-5C hybridoma ^44^), or anti-VTA1 (1:1500, Invitrogen, PA5-21831) for 16 h at 4 °C, followed by peroxidase goat anti-mouse IgG (1:500, Jackson, Cat 115-035-166) or peroxidase goat anti-rabbit IgG (1:500, Jackson, Cat 111-035-144) for 1 hour at room temperature.

## Acknowledgments

We would like to thank Prof. Michael Kozlov for helpful discussions. This work was supported by the Israel Science Foundation Grants 234/17 (Rousso lab) and 1323/18 (Elia lab).

## Notes

### Competing Interest Statement

The authors have declared no competing interest.

